# Whole Exome Sequencing in 20,197 Persons for Rare Variants in Alzheimer Disease

**DOI:** 10.1101/305631

**Authors:** Neha S. Raghavan, Adam M. Brickman, Howard Andrews, Jennifer J. Manly, Nicole Schupf, Rafael Lantigua, The Alzheimer’s Disease Sequencing Project, Charles J. Wolock, Sitharthan Kamalakaran, Slave Petrovski, Giuseppe Tosto, Badri N. Vardarajan, David B. Goldstein, Richard Mayeux

## Abstract

**Objective:** The genetic bases of Alzheimer’s disease remain uncertain. An international effort to fully articulate genetic risks and protective factors is underway with the hope of identifying potential therapeutic targets and preventive strategies. The goal here was to identify and characterize the frequency and impact of rare and ultra-rare variants in Alzheimer’s disease using whole exome sequencing in 20,197 individuals.

**Methods:** We used a gene-based collapsing analysis of loss-of-function ultra-rare variants in a case-control study design with data from the Washington Heights-Inwood Columbia Aging Project, the Alzheimer’s Disease Sequencing Project and unrelated individuals from the Institute of Genomic Medicine at Columbia University.

**Results:** We identified 19 cases carrying extremely rare *SORL1* loss-of-function variants among a collection of 6,965 cases and a single loss-of-function variant among 13,252 controls (p = 2.17 × 10^-8^; OR 36.2 [95%CI 5.8 – 1493.0]). Age-at-onset was seven years earlier for patients with *SORL1* qualifying variant compared with non-carriers. No other gene attained a study-wide level of statistical significance, but multiple top-ranked genes, including *GRID2IP, WDR76* and *GRN*, were among candidates for follow-up studies.

**Interpretation:** This study implicates ultra-rare, loss-of-function variants in *SORL1* as a significant genetic risk factor for Alzheimer’s disease and provides a comprehensive dataset comparing the burden of rare variation in nearly all human genes in Alzheimer’s disease cases and controls. This is the first investigation to establish a genome-wide statistically significant association between multiple extremely rare loss-of-function variants in *SORL1* and Alzheimer’s disease in a large whole-exome study of unrelated cases and controls.

## Introduction

Alzheimer’s disease (AD) is a highly prevalent disorder that dramatically increases in frequency with age, and has no effective treatment or means of prevention. While three causal genes, Amyloid Precursor Protein (*APP*), Presenilin 1 and 2 (*PSEN1* and *PSEN2*), have been established for early-onset AD (age of onset <65 years of age), the rest of the heritability is still unknown. Further, beyond Apolipoprotein E (*APOE*), which confers the greatest risk for late-onset AD (age of onset ≥65 years of age), there remains a large gap in the understanding of its causes. Identifying genetic variants that increase risk or protect against AD is considered an international imperative because of the potential therapeutic targets that may be revealed. Recent technological advances in genome-wide association studies and high throughput next-generation sequencing may help to implicate variants in genes in specific molecular pathways relevant to AD.

In this study, we used whole-exome sequencing to investigate all protein-coding genes in the genome focusing on ultra-rare (allele frequency less than 0.01%) and putatively deleterious variants. Rare variants are hypothesized to contribute to disease ^1, 2^, and studies of complex traits in population genetic models indicate an inverse relationship between the odds ratio and effect size conferred by rare variants and low allele frequencies ^3^. Thus, we searched for large effects conferred by putatively causal ultra-rare variants. Traditional single variant statistics can be underpowered because patients with similar clinical presentations possess distinct rare variants that inflict similar effects on the gene ^4^. Gene-based collapsing analyses increase signal detection by aggregating individual qualifying variants within an *a priori* region (e.g., a gene), facilitating detection of genes associated with disease through a specific class of genetic variation (e.g., loss-of-function variants).

In order to maximize the ability to detect ultra-rare variants associated with AD, exome-sequencing data of 20,197 cases and controls from the Washington Heights-Inwood Community Aging Project (WHICAP), the Alzheimer’s Disease Sequencing Project (ADSP) and unrelated controls from the Institute of Genomic Medicine were systematically combined and analyzed using a collapsing method with proven prior success in identifying disease associated genes ^5, 6^.

## Methods

The three groups used in this study and their sequencing information are described below.

### Washington Heights-Inwood Community Aging Project

The WHICAP study consisted of a multi-ethnic cohort of 4,100 individuals followed over several years The cohort participants were non-demented initially, 65 years of age or older, and comprised of non-Hispanic whites, African Americans, and Caribbean Hispanics from the Dominican Republic. During each assessment, participants received a neuropsychological test battery, medical interview, and were reconsented for sharing of genetic information and autopsy. A consensus diagnosis was derived for each participant by experienced clinicians based on NINCDS-ADRDA criteria for possible, probable, or definite AD, or moderate or high likelihood of neuropathological criteria of AD ^7, 8^. Every individual with whole-exome sequencing has at least a baseline and one follow-up assessment and examination, and for those who have died, the presence or absence of dementia was determined using a brief, validated telephone interview with participant informants: the Dementia Questionnaire (DQ) ^9^ and the Telephone Interview of Cognitive Status (TICS) ^10^. 3,702 exome-sequenced WHICAP individuals were designated with case or control status and included in this analysis. From the sequenced cohort, 27% died and less than 1% were lost at follow-up.

### Alzheimer’s Disease Sequencing Project

The ADSP, developed by the National Institute on Aging (NIA) and National Human Genome Research Institute (NHGRI) includes a large case-control cohort of approximately 10,000 individuals ^7^. The recruitment of these individuals was in collaboration with the Alzheimer’s Disease Genetics Consortium and the Cohorts for Heart and Aging Research in Genomic Epidemiology Consortium. The details and rationale for the case-control selection process have been previously described ^7^. All cases and controls were at least 60 years old and were chosen based on sex, age and *APOE* status: 1) controls were evaluated for their underlying risk for AD and for their likelihood of conversion to AD by age 85, based on age at last examination, sex, and *APOE* genotype, and those with the least risk for conversion to AD were selected, and 2) cases were evaluated for their underlying risk for AD based on age at onset, sex, and *APOE* genotype and those with a diagnosis least explained by these factors were selected ^7^. Cases were determined either because they met NINCDS-ADRDA clinical criteria for AD, or postmortem findings met moderate or high likelihood of neuropathological criteria of AD ^^7, 8^^. Autopsy data was available for 28.7% of the cases and controls used in the analysis. Further, some cases were originally diagnosed clinically, subsequently died and had neuropathological findings available after postmortem examination. Cases had documented age at onset or age at death (for pathologically determined cases). Controls were free of dementia by direct, documented cognitive assessment or neuropathological results. The ADSP group consisted of European-Americans and Caribbean Hispanics. All data were available for download for approved investigators at The National Institute on Aging Genetics of Alzheimer’s Disease Data Storage Site website (https://www.niagads.org/adsp/content/home). As part of the ADSP, 116 non-Hispanic white WHICAP controls and 34 cases previously sequenced were included here.

### Additional Controls

The Institute for Genomic Medicine (IGM) (Columbia University Medical Center, New York, NY) hosts an internal database of sequencing data collected from previously exome-sequenced material. In this study, exome-sequencing data from 6,395 IGM controls were utilized. All data used were previously consented for future control use from multiple studies of various phenotypes. The cohort was made up of 55.7% healthy controls and 46.3% with diseases not co-morbid with AD (disease classifications shown in Supplemental Table 1). Although the cohort of controls were not enriched for any neurological disorder or diseases with a known co-morbidity with AD, presence or future possibility of AD could not be excluded based on the available clinical data. individuals with Age and *APOE* status were not available for these participants. The cohort comprised of 70% non-Hispanic white individuals along with those of African American, Hispanic, Middle Eastern, Asian and unknown descent.

### Sequencing, Quality Control and Variant Calling

Whole-exome sequencing of the WHICAP cohort was performed at Columbia University. The additional controls were sequenced at Duke University and Columbia University. Whole-exome sequencing of the ADSP cohort was performed at The Human Genome Sequencing Center, Baylor College of Medicine, Houston, Texas; The Broad Institute Sequencing Platform, The Eli & Edythe L. Broad Institute of the Massachusetts Institute of Technology and Harvard University, Cambridge Massachusetts and Washington University Genome Sequencing Center, Washington University School of Medicine, Saint Louis, Missouri. ADSP raw files in the sequencing read archive format were downloaded from the dbGAP database and decompressed to obtain FASTQ files.

All data were reprocessed for a consistent alignment and variant calling pipeline consisting of the primary alignment and duplicate marking using the Dynamic Read Analysis for Genomics (DRAGEN) platform followed by variant calling according to best practices outlined in Genome Analysis Tool Kit (GATK v3.6). Briefly, aligned reads were processed for indel realignment followed by base quality recalibration and Haplotype calling to generate variant calls. Variant calls were then subject to Variant Quality Score Recalibratrion (VQSR) using the known single nucleotide variants (SNVs) sites from HapMap v3.3, dbSNP, and the Omni chip array from the 1000 Genomes Project. SNVs were required to achieve a tranche of 99.9% and indels a tranche of 95%. Finally, read-backed phasing was performed to determine phased SNVs and merge multinucleotide variants (MNVs) when appropriate. Variants were annotated using ClinEff with Ensembl-GRCh37.73 annotations.

Quality thresholds were set based on previous work^5, 6^, such that all resulting exome variants had a quality score of at least 50, quality by depth score of at least 2, genotype quality score of at least 20, read position rank sum of at least −3, mapping quality score of at least 40, mapping quality rank sum greater than −10, and a minimum coverage of at least 10. SNVs had a maximum Fisher’s strand bias of 60, while indels had a maximum of 200. For heterozygous genotypes, the alternative allele ratio was required to be greater than or equal to 25% and variant from sequencing artifacts and exome variant server failures (http://evs.gs.washington.edu/EVS) were excluded.

Quality control was performed on all sequencing data. Samples with less than 90% of the consensus coding sequence (CCDS) covered at 10X and samples with sex-discordance between clinical and genetic data were excluded from the analysis. Cryptic relatedness testing was performed using KING, and second degree or closer (relatedness threshold of 0.0884 or greater) relatives were removed with preferential retention of cases over controls and subsequently samples with higher average read-depth coverage.

The consensus coding sequence ^11^ (CCDS) annotated protein-coding region for each gene (n=18,834) was tabulated as either carrying or not carrying a qualifying variant for every individual. Qualifying variants were defined for a loss-of-function model: stop gain, frameshift, splice site acceptor, splice site donor, start lost, or exon deleted variants. A negative control analysis was performed defining qualifying variants as synonymous variants to detect potential biases in variant calling between the cases and controls separately for each of the top four genes. The minor allele frequency threshold was 0.01% internally and within African American, Latino and Non-Finnish European populations from the Exome Aggregation Consortium^12^ (ExAC release version 0.3.1). The allele frequency thresholds use a “leave-one-out” method for the combined test cohort of cases and controls such that the minor allele frequency of each variant was calculated using all individuals except for the index sample under investigation. Thus, the maximum instances of a single variant a gene in our sample of 20,197 was five. A dominant model was defined such that one or more qualifying variant(s) in a gene qualified the gene.

An important aspect of the collapsing analysis methodology is the reduction of variant calling bias due to coverage differences between cases and controls. To ensure balanced sequencing coverage of evaluated sites between cases and controls, we imposed a statistical test of independence between the case/control status and coverage. For a given site, consider ***s*** total number of cases, ***t*** total number of controls and ***x*** number of cases covered at 10X, ***y*** number of controls covered at 10x. We model the number of covered cases X as a Binomial random variable:

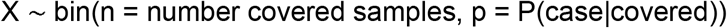

If case/control status and coverage status are independent, then:

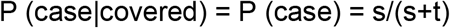

We can test for this independence by performing a two-sided Binomial test on the number of covered samples at given site, x.

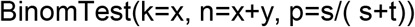

In the collapsing analyses, a binomial test for coverage balance as described above was completed as an additional qualifying criterion. Any site which resulted in a nominal significance threshold of 0.05 was eliminated from further consideration.

A Fisher’s exact test on qualifying variants in cases and controls for each gene was performed and imbalances in cases and controls within a gene indicated a possible association with the case-ascertained phenotype. Ultra-rare variant analyses were conducted using Analysis Tools for Annotated Variants (ATAV), developed and maintained by the Institute for Genomic Medicine at Columbia University. Study-wise significance was set to 0.05/18,834(# of genes tested) = 2.7×10^-6^. Fisher’s Exact Test for the polygenic comparison of International Genetics of Alzheimer’s Project (IGAP) loci ^13^ and t-test for age of onset-analysis (presented as mean +/-standard deviation) were conducted in R v.3.3.1.

## Results

We analyzed the exomes of 6,965 individuals meeting with the diagnosis of AD and 13,232 controls (Table 1). Prior to analysis, 570 individuals (91 cases and 479 controls) were removed due to known or cryptic relatedness. For ultra-rare variant analysis (MAF of 0.01% or lower), conventional population stratification has not been a strong confounder as it can be in common variant analyses; and these results did not significantly differ from meta-analyses in population stratified data. All variants reported here were found in five or less individuals from the study, and most variants were found in only one person, increasing the confidence that population stratification was not an issue. An important distinction exists between the cases and controls in the ADSP and WHICAP datasets. In the ADSP dataset, the younger cases were preferentially chosen as part of the study design ^7^. The WHICAP individuals are part of a population-based cohort followed longitudinally, and thus cases were older than controls.

**Table 1.**
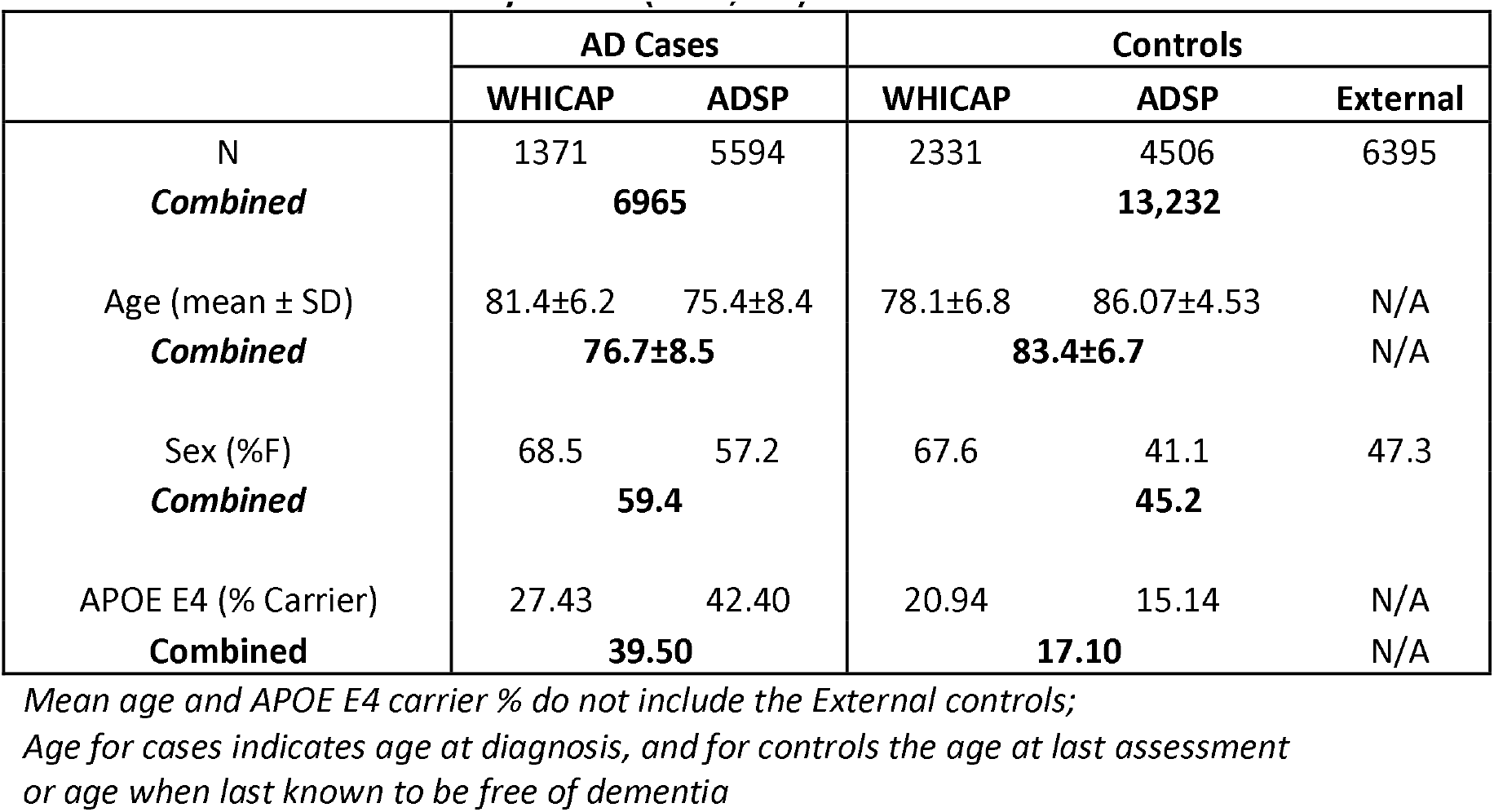
Characteristics of Study Cohort (n=20,197)

Of the 18,834 genes analyzed, 15,736 contained at least one qualifying variant. Genomic inflation for the analysis was very modest, λ = 1.04 (Figure 1). Gene-based, collapsing analyses for loss-of-function variants, with allele frequency less than 0.01% (within the study cohort, and separately within ExAC^12^) identified *SORL1* to be enriched in cases compared to controls at an exome-wide significance level of p = 2.17 × 10^-8^ (Table 2). We confirmed the results for *SORL1* were not driven by a particular ethnicity by running individual association tests on non-Hispanic Whites, Caribbean Hispanics, and African Americans as described above, separately and summarizing them in a sample weight meta-analysis^14^*(SORL1* p = 2.45 × 10^-8^). Although no other gene attained the study-wide level of statistical significance, *GRID2IP* (p = 2.98 × 10^-4^), *WDR76* (p = 7.39 × 10^-4^) and *GRN* (p = 9.56 × 10^-4^) were highly-ranked candidate genes that were case-enriched for loss-of-function variants (Table 2). Extended results are found in **Supplemental Table 2**. There were no significant differences in synonymous variation in these four genes (1.5% cases, 1.7% of controls; FET p = 0.25).

**Figure 1.**
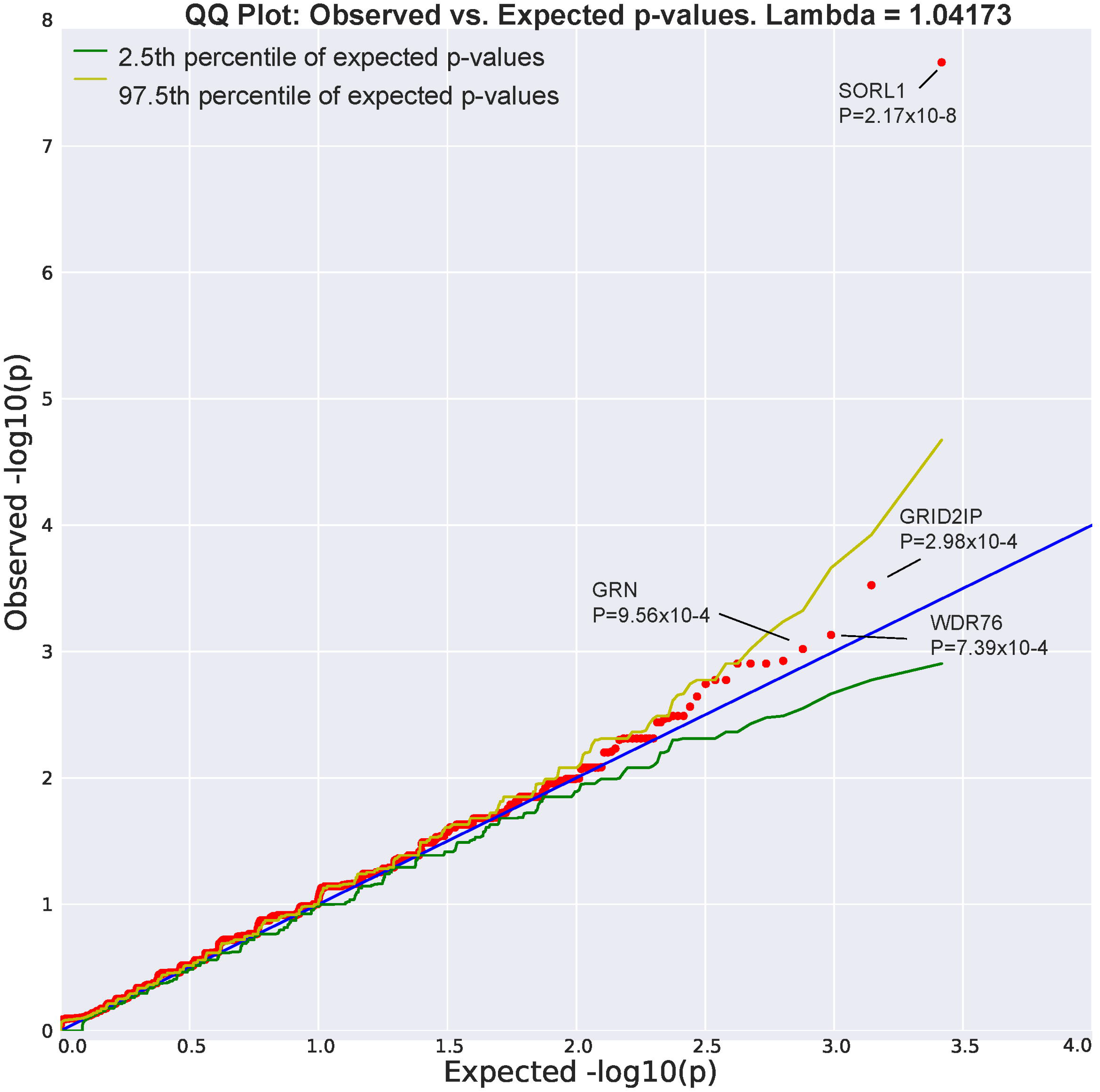
QQ Plot: Observed vs. expected p-values. Lambda = 1.04173

**Table 2.**
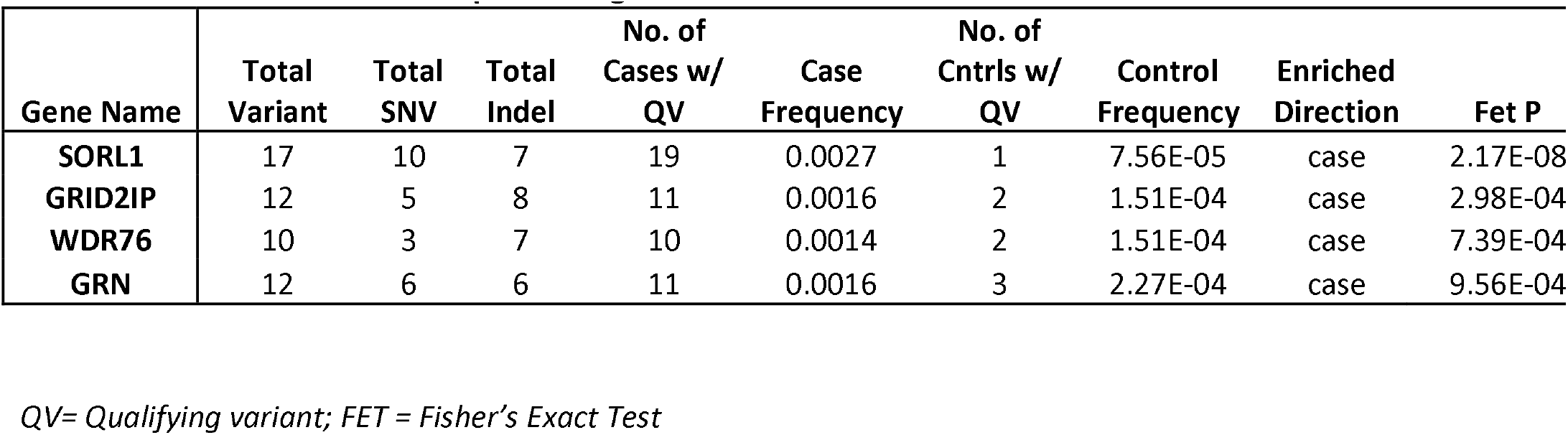
Variant counts for the top four AD genes

There were 19 cases with a loss-of-function qualifying variant in *SORL1*(Table 3) among 6,965 cases (frequency = 0.27%) and one variant among 13,232 controls (frequency = 0.0076%). Given the rate of *SORL1* loss-of-function qualifying variants found in our control sample (1 / 13,232; frequency = 0.0076%), we expected to identify only 0.5 loss-of-function variants by chance among our 6,965 cases; however, we identified 19. The accompanying odds ratio for AD risk upon identifying a *SORL1* loss-of-function qualifying variants as defined in this study was 36 [95% CI 5.8 – 1493.0]. Targeted investigation into the single control indicated a diagnosis of mild cognitive impairment^15^. The *SORL1* loss-of-function variants were found across the non-Hispanic white, Caribbean Hispanic, and African American cases. Six of the 19 cases were deceased with autopsy confirmation of the AD diagnosis^16^.

**Table 3.** SORL1 variants

| Genomic Position | Variant Type | Variant Class | CADD score | Protein modification | ExAC Global Frequency | Case/Control | Sex | Ethnicity | Braak Stage | Age at Onset or Last Visit |
| --- | --- | --- | --- | --- | --- | --- | --- | --- | --- | --- |
| 11-121367577 | snv | SAV | 26.6 | NA | 0 | case | F | AA | NA | 77 |
| 11-121367654 | snv | SG | 37 | p.Arg279* | 0 | case | F | NHW | 6 | 72 |
| **11-12142134322<sup>23</sup>** | snv | SG | 39 | p.Arg744* | 0 | case | M | NHW | NA | 65 |
| **11-12142134322<sup>23</sup>** | snv | SG | 39 | p.Arg744* | 0 | case | F | NHW | NA | 67 |
| 11-121426001 | indel | FV | NA | p.Asp850fs | 0 | case | F | NHW | NA | 60 |
| 11-121428047 | snv | SG | 41 | p.Arg866* | 0 | case | M | NHW | 6 | 65 |
| 11-121430263 | indel | FV | NA | p.Ile983fs | 0 | ctrl | M | AA | NA | 64 |
| 11-121440980 | snv | SDV | 27.6 | NA | 4.95E-05 | case | F | CH | NA | 80 |
| 11-121456930 | snv | SAV | 26.8 | NA | 0 | case | M | NHW | NA | 69 |
| 11-121456930 | snv | SAV | 26.8 | NA | 0 | case | M | NHW | 6 | 62 |
| 11-121461788 | indel | FV | NA | p.Cys1431fs | 0 | case | F | NHW | NA | 61 |
| **11-12146648224<sup>25</sup>** | snv | SDV | 28 | NA | 0 | case | F | NHW | 3 | 90+ |
| **11-12146648224<sup>25</sup>** | snv | SDV | 28 | NA | 0 | case | F | NHW | NA | 90+ |
| 11-121474911 | indel | FV | NA | p.Thr1511fs | 0 | case | M | NHW | NA | 60 |
| 11-121474984 | snv | SG | 35 | p.Cys1534* | 0 | case | F | NHW | NA | 74 |
| **11-12147756824<sup>25</sup>** | snv | SG | 46 | p.Arg1655* | 0 | case | M | NHW | NA | 69 |
| 11-121477667 | snv | SDV | 26.9 | NA | 0 | case | F | AA | NA | 68 |
| 11-121485637 | indel | FV | NA | p.Asp1828fs | 0 | case | M | NHW | NA | 75 |
| 11-121491801 | indel | FV | NA | p.Lys1975fs | 0 | case | M | NHW | 6 | 61 |
| 11-121500253 | indel | FV | NA | p.Met2211fs | 0 | case | M | NHW | 6 | 62 |

Those in bold have previously been identified as indicated by the reference
SNV = Single Nucleotide Variant; Indel = Insertion or Deletion
CADD = Combined Annotation Dependent Depletion
FV = Frameshift Variant; SAV = Splice Acceptor Variant; SDV = Splice Donor Variant; SG = Stop Gained
AA = African American; CH = Caribbean Hispanic; NHW = Non-hispanic White

Of relevance to loss-of-function variant case-enrichment, *SORL1* is known to be among the protein-coding genes most significantly depleted of loss-of-function variants in the general population (LOF depletion FDR = 2 × 10^-7^) (Table 2). Of the 17 distinct *SORL1* loss-of-function qualifying variants, only one (11:121440980, rs200504189) was found in the ExAC database^12^. *SORL1* was also significantly enriched for functional variants (nonsynonymous and predicted as possibly or probably damaging by PolyPhen-2 HumVar^17^) (p = 9.79×10^-7^), 1.8% of cases had a qualifying functional variant compared to 1% controls. There was no difference in the frequency of *APOE-ε4* carriers among cases with qualifying variants in *SORL1* compared to those without these variants (40.0% vs. 39.6%). Age-at-onset analyses revealed a 6.81 year difference between cases with a *SORL1* qualifying variant versus non-carrying cases (AD carriers: 69.86 +/-9.37; AD non-carriers: 76.67 +/-8.53; t(6963), p = 4x10^-4^).

Coverage for the 12 qualifying *GRID2IP* variants was lower in the sequencing performed in this project and in ExAC^12^, reducing our confidence of the rare variant calling for this gene because it is likely not represented well by exome capture libraries. The median of mean read-depth coverage of the *GRID2IP* variants was 21-fold and at these exact same sites in ExAC^12^, 4-fold. However, read-depth coverage was higher in the genome aggregation database (gnomAD), with a median of mean read-depth coverage of 21-fold, and only two loss-of-function variants less than the 0.0001 allele frequency threshold. Two of the 11 cases were deceased with autopsy confirming the pathological diagnosis of AD^16^.

Coverage for *WDR76* and *GRN* were excellent in this study and in ExAC^12^. Three of the 10 individuals clinically diagnosed as AD with loss-of-function qualifying variants in *WDR76* had undergone autopsy. One met postmortem criteria defined as high likelihood of Alzheimer’s disease, a second met intermediate likelihood^16^, however, the third had no distinctive pathology and no definitive diagnosis was derived. Two of the 11 individuals with *GRN* loss-of-function qualifying variants had autopsy data; one met criteria for AD and the other for frontotemporal lobar degeneration (FTLD) ^18^. None of the GRN carriers carried variants in any of the top four genes.

We also investigated rare variants in loci that were associated with AD in the IGAP genome wide association study ^13^ along with *APP, PSEN1, PSEN2*, and *TREM2.*(Table 4). Qualifying variants in *SORL1* and *ZCWPW1* (p=0.02) were more frequent in cases than controls. Overall, there was a slight increase in the frequency of variants in cases compared with controls (Fisher’s exact p=0.002), but after the removal of *SORL1*, the association was no longer significant (Fisher’s exact p=0.11).

**Table 4.**
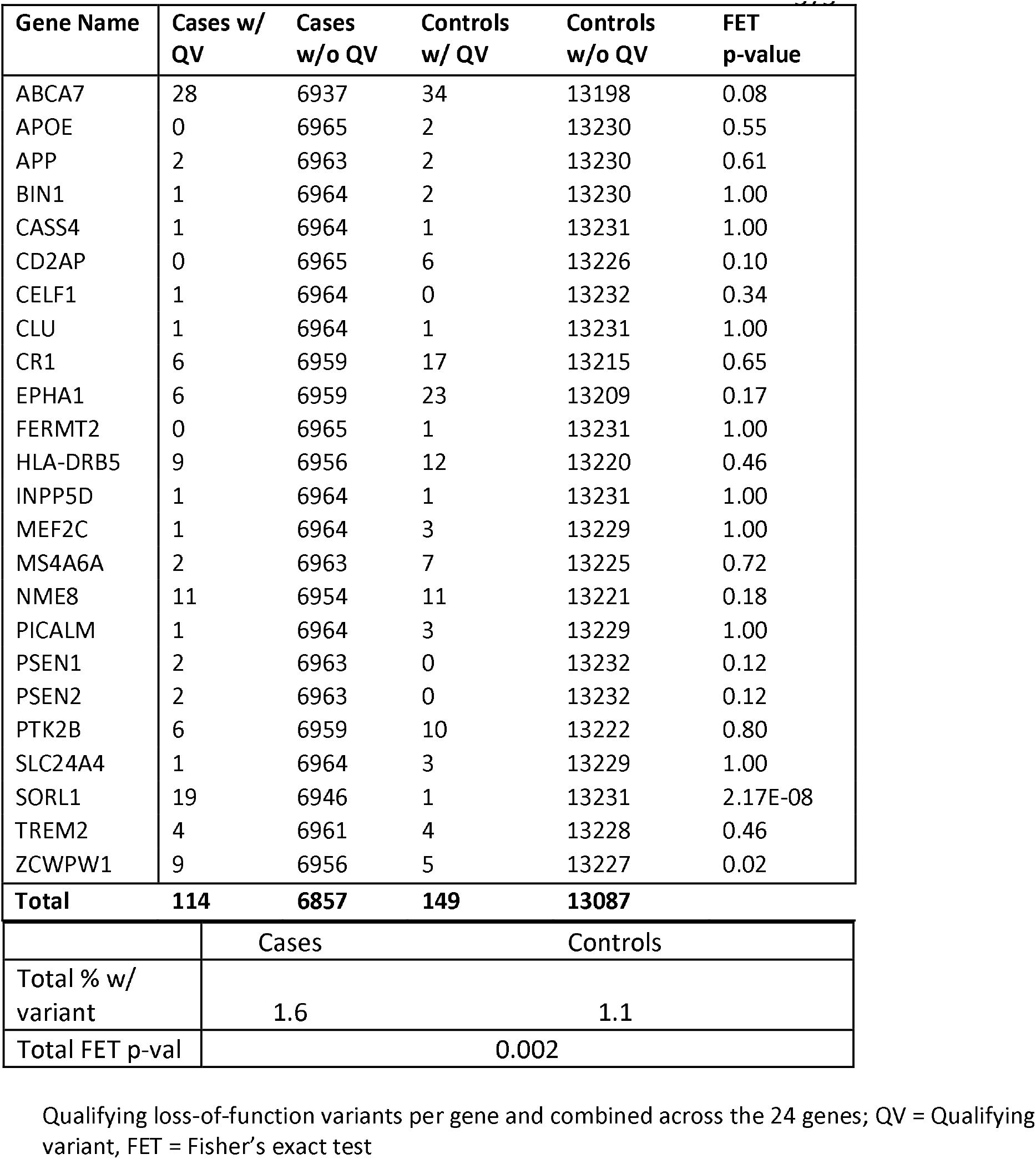
Counts of ultra-rare variant in previously identified or implicated AD genes

## Discussion

This study provides strong evidence that ultra-rare, loss-of-function variants in *SORL1* represent an important genetic risk factor for AD. This is the first investigation to establish a genome-wide statistically significant association between ultra-rare variants in *SORL1* and AD in a large, unbiased whole-exome study of unrelated early- and late-onset cases and controls. *SORL1* has previously been implicated in both familial and sporadic, early- and late-onset Alzheimer’s disease ^19–25^.

Common variants in *SORL1* were first genetically associated with AD in a candidate gene analysis using 29 common variants ^24^. Shortly thereafter, nine rare loss-of-function variants including nonsense, frameshift and splice site mutations were described in familial and sporadic early onset AD ^19, 20^. The *SORL1* findings in early onset AD were replicated in larger European cohorts of patients^21^. Using a targeted, candidate gene approach, *SORL1* variants were found by us in familial and sporadic late-onset AD among Caribbean Hispanics as well as patients with European ancestry with sporadic late-onset AD ^26^. Our findings here indicated that cases who possess a *SORL1* qualifying variant were on average younger at onset. Yet, only four of the cases with a *SORL1* qualifying variant were diagnosed before the age of 65, implicating that the gene is involved in both early- and late-onset AD.

Holstege, et al. ^23^, reported that strongly damaging, but rare variants (E×AC^12^ MAF < 1×10^-5^) in *SORL1* as defined by a Combined Annotation Dependent Depletion (CADD) score of greater than 30, increased the risk of Alzheimer’s disease by 12-fold. The authors proposed that the presence of these variants should be considered in addition to risk variants in *APOE*, and causal variants in *PSEN1, PSEN2* or *APP* for assessing risk in a clinical setting. Accordingly, only one of the *SORL1* variants identified in our study was found in ExAC^12^, and was very rare (11:121440980; E×AC AF = 4.95×10^-5^). Furthermore, half of the 10 variants with a CADD score available were over 30, and all were over 25. The depletion of loss-of-function variants in the ExAC database lends further evidence to the significance of the higher frequency of loss-of-function variants in our AD sample.

*SORL1*, also known as *SORLA* and *LR11*, encodes a trafficking protein (sortilin-related receptor, L(DLR class) A repeats containing protein) that binds the amyloid precursor protein (APP) redirecting it to a non-amyloidogenic pathway within the retromer complex. The major site for expression of SORL1 protein is in the brain especially within neurons and astrocytes. Aβ peptides are also directed to the lysosome by SORL1. Processing of APP requires endocytosis of molecules from the cell surface to endosomes whereby proteolytic breakdown to Aβ occurs. SORL1 acts as a sorting receptor for APP that recycles molecules from endosomes back to the trans-Golgi network to decrease Aβ production. We found that in the absence of the SORL1 gene, APP was released into the late endosome where it underwent β-secretase and γ- secretase cleavage generating Aβ ^24^. Thus, the mechanisms by which mutations in SORL1 lead to neurodegeration in Alzheimer’s disease relates to the disruption of its ability to bind APP. Qualifying variants in other genes were also more prevalent among patients with AD compared with healthy, non-demented controls. Variants in *GRID2IP, WDR76* and *GRN* were four to five times more frequent in cases than in controls, though these genes have not yet achieved genome-wide significance and thus further studies including larger patient samples will help determine which contribute to AD risk.

Glutamate receptor delta-2 interacting protein (*GRID2IP*) is selectively expressed in the cerebellar Purkinje cell-fiber synapses. The exact role for this gene is not fully understood, but it appears to be a postsynaptic scaffold protein that links to GRID2 with signaling molecules and the actin cytoskeleton ^27^. There is no known role for *GRID2IP* in AD despite the fact that mutations were found in two individuals with postmortem confirmed Alzheimer’s disease. The gene has not been well represented in existing exome sequencing libraries and the resulting reduced coverage of this gene makes the findings more difficult to interpret. However, the variants driving the signal in our analyses are all well covered in our entire cohort, with more than 96% of samples achieving at least 10X coverage.

*WDR76* interacts with chromatin components and the cytosolic chaperonin containing TCP-1 (CCT), allowing for the maintenance of cellular homeostasis by assisting in the identification of folded proteins. *WDR76* has low expression in brain and relatively high expression in lymph nodes. Only one of the three individuals with postmortem data met “high likelihood criteria” for AD.

*GRN* mutations in patients with clinically diagnosed AD have been previously reported in large families in the National Institute on Aging family-based study (NIA-AD) ^28^ and among large, multiply affected families of Caribbean Hispanic ancestry ^29^. These loss-of-function mutations result in haploinsufficiency, premature stop codons or nonsense variants impairing the secretion or the structure of Progranulin, involved intracellular trafficking and lysosomal biogenesis and function. Its role in AD in unclear and possibly coincidental ^30^. The phenotype of FTLD includes unique manifestations allowing it to be distinguished from AD. A family presumed to have Alzheimer’s disease phenotypically with a *GRN* mutation (c.154delA) had FTLD with ubiquitin-positive, tau-negative and lentiform neuronal intranuclear inclusions (-U NII) with neuronal loss and gliosis affecting the frontal and temporal lobes, and TDP43 inclusions ^31^. Only one of the six family members (Patient II:1) had mixed pathology meeting NIA-Reagan criteria of high likelihood ^16^ and coexisting FTLD-U N11 with TDP43 inclusions. *GRN* mutations were also observed in a sporadic patient with postmortem evidence of Alzheimer’s disease: NIA-Reagan criteria of high likelihood^16^ and coexisting FTLD-U N11 with TDP43 inclusions ^32^. Among the patients with *GRN* mutations in this study, one patient met criteria for definite Alzheimer’s disease without co-existing FTLD, while another met pathological criteria for FTLD.

The results here indicate that extremely rare, loss-of-function variants in *SORL1* have an strongly effect the risk of sporadic AD. While qualifying variants were present in only 0.27% of patients, only a single variant was found among 13,232 controls, and the single control carrier upon a post hoc cognitive evaluation was identified to have a diagnosis of mild cognitive impairment. These results confirm and greatly extend those from sequencing studies in familial and sporadic early onset Alzheimer’s disease ^19–21^, familial AD families ^24, 26, 33^ and investigations within clinical settings. The resulting impact of the loss-of-function variants in *SORL1* on recycling of the amyloid precursor protein and the amyloid β protein make this pathway an attractive target for the development of therapies. Beyond implicating *SORL1* and highly suggestive candidate genes for AD, this study shows for the first time that the collapsing analysis methodology of ultra-rare variants described here that has proven successful for a number of rare diseases also can securely implicate genes in a condition as common as AD.

## Author Contributions

### Study Design

NSR, CW, SK, SP, GT, BNV, DBG, RM

### Data Collection

AMB, HA, JJM, NS, RL, CW, SK, SP, GT, BNV, DBG,

### RMData Analysis

NSR, CW, SK, SP, GT, BNV, DBG, RM

### Writing and Editing

NSR, AMB, HA, JJM, NS, RL, CW, SK, SP, GT, BNV, DBG, RM

## Acknowledgements

### WHICAP and EFIGA

Data collection for this project was supported by the Washington Heights and Inwood Community Aging Project (WHICAP) and Genetic Studies of Alzheimer’s disease in Caribbean Hispanics (Estudio familiar de la genètica de la enfermedad de Alzheimer, also known as EFIGA) funded by the National Institute on Aging (NIA), by the National Institutes of Health (NIH) (1RF1AG054023, 5R37AG015473, RF1AG015473, R56AG051876), and the National Center for Advancing Translational Sciences, NIH through Grant Number TL1TR001875. We acknowledge the WHICAP and EFIGA study participants and the research and support staff for their contributions to this study.

### ADSP

The Alzheimer’s Disease Sequencing Project (ADSP) is comprised of two Alzheimer’s Disease (AD) genetics consortia and three National Human Genome Research Institute (NHGRI) funded Large Scale Sequencing and Analysis Centers (LSAC). The two AD genetics consortia are the Alzheimer’s Disease Genetics Consortium (ADGC) funded by NIA (U01 AG032984), and the Cohorts for Heart and Aging Research in Genomic Epidemiology (CHARGE) funded by NIA (R01 AG033193), the National Heart, Lung, and Blood Institute (NHLBI), other National Institute of Health (NIH) institutes and other foreign governmental and non-governmental organizations. The Discovery Phase analysis of sequence data is supported through UF1AG047133 (to Drs. Schellenberg, Farrer, Pericak-Vance, Mayeux, and Haines); U01AG049505 to Dr. Seshadri; U01AG049506 to Dr. Boerwinkle; U01AG049507 to Dr. Wjsman; and U01AG049508 to Dr. Goate and the Discovery Extension Phase analysis is supported through U01AG052411 to Dr. Goate and U01AG052410 to Dr. Pericak-Vance. Data generation and harmonization in the Follow-up Phases is supported by U54AG052427 (to Drs. Schellenberg and Wang).

The ADGC cohorts include: Adult Changes in Thought (ACT), the Alzheimer’s Disease Centers (ADC), the Chicago Health and Aging Project (CHAP), the Memory and Aging Project (MAP), Mayo Clinic (MAYO), Mayo Parkinson’s Disease controls, University of Miami, the Multi-Institutional Research in Alzheimer’s Genetic Epidemiology Study (MIRAGE), the National Cell Repository for Alzheimer’s Disease (NCRAD), the National Institute on Aging Late Onset Alzheimer’s Disease Family Study (NIA-AD; U24 AG056270), the Religious Orders Study (ROS), the Texas Alzheimer’s Research and Care Consortium (TARC), Vanderbilt University/Case Western Reserve University (VAN/CWRU), the Washington Heights-Inwood Columbia Aging Project (WHICAP) and the Washington University Sequencing Project (WUSP), the Columbia University Hispanic-Estudio Familiar de Influencia Genetica de Alzheimer (EFIGA), the University of Toronto (UT), and Genetic Differences (GD).

The CHARGE cohorts, with funding provided by 5RC2HL102419 and HL105756, include the following: Atherosclerosis Risk in Communities (ARIC) Study which is carried out as a collaborative study supported by NHLBI contracts (HHSN268201100005C, HHSN268201100006C, HHSN268201100007C, HHSN268201100008C, HHSN268201100009C, HHSN268201100010C, HHSN268201100011C, and HHSN268201100012C), Austrian Stroke Prevention Study (ASPS), Cardiovascular Health Study (CHS), Erasmus Rucphen Family Study (ERF), Framingham Heart Study (FHS), and Rotterdam Study (RS). CHS research was supported by contracts HHSN268201200036C, HHSN268200800007C, N01HC55222, N01HC85079, N01HC85080, N01HC85081, N01HC85082, N01HC85083, N01HC85086, and grants U01HL080295 and U01HL130114 from the National Heart, Lung, and Blood Institute (NHLBI), with additional contribution from the National Institute of Neurological Disorders and Stroke (NINDS). Additional support was provided by R01AG023629, R01AG15928, and R01AG20098 from the National Institute on Aging (NIA). A full list of principal CHS investigators and institutions can be found at CHS-NHLBI.org. The content is solely the responsibility of the authors and does not necessarily represent the official views of the National Institutes of Health. The three LSACs are: the Human Genome Sequencing Center at the Baylor College of Medicine (U54 HG003273), the Broad Institute Genome Center (U54HG003067), and the Washington University Genome Institute (U54HG003079).

Biological samples and associated phenotypic data used in primary data analyses were stored at Study Investigators institutions,and at the National Cell Repository for Alzheimer’s Disease (NCRAD, U24AG021886) at Indiana University funded by NIA. Associated Phenotypic Data used in primary and secondary data analyses were provided by Study Investigators, the NIA funded Alzheimer’s Disease Centers (ADCs), and the National Alzheimer’s Coordinating Center (NACC, U01AG016976) and the National Institute on Aging Genetics of Alzheimer’s Disease Data Storage Site (NIAGADS, U24AG041689) at the University of Pennsylvania, funded by NIA, and at the Database for Genotypes and Phenotypes (dbGaP) funded by NIH. This research was supported in part by the Intramural Research Program of the National Institutes of health, National Library of Medicine. Contributors to the Genetic Analysis Data included Study Investigators on projects that were individually funded by NIA, and other NIH institutes, and by private U.S. organizations, or foreign governmental or nongovernmental organizations.

We would like to acknowledge the following individuals or groups for the contributions of control samples: T. Young; K. Whisenhunt; S. Palmer; S. Berkovic, I. Scheffer, B. Grinton; E. Cirulli; M. Winn; R.Gbadegesin; A. Poduri; S. Schuman; E. Nading; E. Pras; D. Lancet; Z. Farfel; S. Kerns; H. Oster; D. Valle; J. Hoover-Fong; N. Sobriera; M. Hauser; G. Nestadt; J. Samuels; Y. Wang; G. Cavalleri, N. Delanty; C. Depondt; S. Sisodiya; R. Buckley; C. Moylan; A. M. Diehl; M. Abdelmalek; S. Delaney; V. Shashi; M. Walker; M. Sum; the ALS Sequencing Consortium; the Washington University Neuromuscular Genetics Project; DUHS (Duke University Health System) Nonalcoholic Fatty Liver Disease Research Database and Specimen Repository; Epilepsy Genetics Initiative, A Signature Program of CURE; the Epi4K Consortium and Epilepsy Phenome/Genome Project; the Undiagnosed Diseases Network; and the Utah Foundation for Biomedical Research.

The collection of control samples and data was funded in part by: Biogen; Gilead Sciences, Inc.; UCB; National Institutes of Health (RO1HD048805); National Institute of Neurological Disorders and Stroke (U01NS077303, U01NS053998, U54NS078059); National Institute of Child Health and Human Development (P01HD080642); National Institute of Mental Health (R01MH097971, K01MH098126); National Human Genome Research Institute (U01HG007672); an American Academy of Child and Adolescent Psychiatry (AACAP) Pilot Research Award; Endocrine Fellows Foundation Grant; the NIH Clinical and Translational Science Award Program (UL1TR000040); the Ellison Medical Foundation New Scholar award AG-NS-0441-08; Duke Chancellor’s Discovery Program Research Fund 2014; The J. Willard and Alice S. Marriott Foundation; The Muscular Dystrophy Association; The Nicholas Nunno Foundation; The JDM Fund for Mitochondrial Research; The Arturo Estopinan TK2 Research Fund; the Stanley Institute for Cognitive Genomics at Cold Spring Harbor Laboratory; New York-Presbyterian Hospital; the Columbia University College of Physicians and Surgeons; and the Columbia University Medical Center.

The content is solely the responsibility of the authors and does not necessarily represent the official views of the National Institutes of Health.

Biogen Inc. provided support for whole exome sequencing for the WHICAP cohort through a grant to David Goldstein, PhD and salary support for Neha S. Raghavan PhD for analyses. Individuals at Biogen were not involved in the collection of data, analysis or interpretation of the genetic data, nor in the production of this manuscript.

